# A look at the ground: time lapse camera trapping to determine the composition and abundance of small sized wildlife communities

**DOI:** 10.1101/2025.08.12.669913

**Authors:** V. García-López, J. Vicente, F. Carro, P. Acevedo, A. Ertürk

## Abstract

Micromammals are critical species in the context of ecosystem functioning, assuming a significant role as prey species and environmental engineers. Nevertheless, they present a challenge when developing monitoring programs. As a proof of concept, a time-lapse based camera-trapping non-invasive protocol was tested with the application of the Space to Event Model (STE) was determined. The objective of this study was to estimate the density of micromammals in Doñana National Park (DNP, Southwest Spain) by comparing estimates against independent ones obtained from capture-mark-recapture (CRM) methodology with a standardized protocol using Sherman traps. The micromammal species richness in the area exhibited a strong correlation with the capture methodology employed. Regarding the estimation of abundance, the densities of micromammals determined by STE were found to be within the range of the estimations of minimum population size determined by CRM. The STE, which was found to be promising, also allowed the determination of the abundance of other small-sized terrestrial wildlife, such as reptiles. Our findings indicate its potential for camera trap multi-species monitoring. In terms of practicability, field and data processing efforts for STE are both viable and will continue to improve as new automatic identification tools are incorporated. It is imperative that the STE protocols are examined in a series of study areas exhibiting varying micromammal densities and species compositions, in addition to other small-sized terrestrial wildlife.

## 1. Introduction

The determination and monitoring of wildlife communities’ composition and population trends, as well as driving factors, facilitates a more profound comprehension of the essential ecological and epidemiological processes, thereby enabling informed decision-making. To achieve this objective, it is essential to develop monitoring tools that are both practical and applicable to neglected or difficult to monitor zoological terrestrial groups (see Moeller et al., 2018; Forsyth et al., 2021; *ENETWILD* consortium et al., 2023). Micromammals are key to terrestrial ecosystems in their capacity as secondary producers, contributing substantially to total animal biomass. They function as the trophic base for numerous predators and may act as important reservoirs for disease. Furthermore, they serve as effective bioindicators of ecological changes, such as the effects of climate or land use change (see Carro et al., 2007; Moreno et al., 2008; 2016; Marfil et al., 2009). The abundance, distribution and population trends of many small mammals and other small-sized species are not well studied (Glen et al., 2013). Consequently, the actual quantitative contribution of these species to animal communities is not fully understood.

The estimation of micromammals’ population size is challenging, primarily due to their elusive and nocturnal nature (Williams et al., 2002). Live capture-recapture methods have been the most frequent, accurate and standardised approach (e. g., Tasker and Dickman 2002; Krebs et al., 2011; Theuerkauf et al., 2011, but they can be unethical and/or impractical as they may harm or kill the animals. The Capture-Mark-Recapture (CMR) or more advanced Spatially Explicit Capture-Recapture (SECR) models are typically employed (Krebs et al., 2011; Ivan et al., 2013; Sun et al., 2014; Gerber and Parmenter, 2015; Romairone et al., 2018). The utilisation of SECR models is favoured due to their capacity to mitigate edge-effect bias and accommodate environmental and biological factors that influence animal movement. Efforts are currently centred on the utilisation of alternative techniques, such as camera-trapping, which obviates the need for live capture (for instance, Nicolau 2020, and Gracanin et al. 2022 provide relevant insights in this regard).

Camera traps (CTs) are non-invasive and potentially efficient sensors, although it has mainly focused on medium-sized to large mammals (Blount et al., 2021; Kleiven et al., 2022). The utilisation of camera trapping facilitates the acquisition of data pertaining to the distribution, behaviour, and population dynamics of animals (e. g., Steenweg et al., 2017). For unmarked species or those non-individually recognisable (e. g., Gilbert et al., 2021; Morin et al., 2022) CT methods presuppose assumptions concerning the movement of animals in relation the sampling process and certain methodologies may necessitate the acquisition of supplementary data, such as detection area, the distance to the detected animal, or the animal movement speed (see Morin et al., 2022).

Recent advancements in CT technology and their deployment have resulted in the expanded use of these techniques to include small mammals (see Nicolau et al., 2020). The utilisation of selfie traps (e. g., Moeller et al., 2018; Gracanin et al., 2022) facilitates the application of the SECR model to both marked individuals (e.g., ears, necessitating physical capture) and all unique individuals identified on camera (e. g., unique facial markings and ear scars, which are frequently unfeasible for small mammal species lacking distinct, recognisable features). The logistical protocols encompass the transport, deployment and removal of such devices in the field. A significant challenge for small-sized wildlife is their low detectability by the passive infrared (PIR) trigger utilised in most CTs. Furthermore, there is a challenge with their identification through photographic evidence (Glein et al., 2013). The utilisation of CT timelapse does not necessitate the deployment of devices to attract and fix wildlife for the purpose of selfies, nor does it rely on motion detection. Moeller et al. (2018) developed a novel methodology for estimating animal density from the encounter rate between animals and CTs, termed the space-to-event (STE) model. The term ’STE’ is defined as the extent of the area sampled until the species is encountered. The STE model necessitates the placement of CTs in a random configuration with respect to animal movement and the collection of samples at a predetermined frequency over time. This approach involves the capture of time-lapse photographs, with the animals themselves not serving as triggers for the CTs. Recent studies have identified STE as a promising and cost-effective method of monitoring unmarked species (see Ausband et al., 2022; Lyte et al., 2023). However, to the best of our knowledge, its utility remains largely unexplored for small wildlife.

For the purpose of determining the density of micromammals, a STE time-lapse camera-trapping protocol was implemented and validated in a selected area of Doñana National Park (DNP, Southwest Spain). The specific objectives of this pilot study were twofold: firstly, to describe a protocol adapted to estimate the density of micromammals (and potentially other small-sized terrestrial wildlife) based on time-lapse camera trapping and the STE method; and secondly, to compare the estimated density from STE with independent information on population abundance in the study area.

## 2. Material and Methods

### 2.1. Study area and populations

The study was conducted in DNP, in the south-eastern part of the Iberian Peninsula (36°59′ N, 6°26′ W, Spain) on the right bank of the Guadalquivir River estuary, which presents a sub-humid Mediterranean climate with Atlantic influence (see Figure 1). DNP is considered to be one of the primary biodiversity hotspots within the Mediterranean Basin, and it is a subject to considerable habitat loss (Moreno and Rouco; 2013Green et al., 2024). The biotypes that characterise this biome include dunes, marshes and scrubland (see Carro et al., 2019; Sereno-Cardieno et al., 2023). The present study was conducted in the scrub biotope of the northern Matasgordas area (Figure 1).

**Figure 1.**
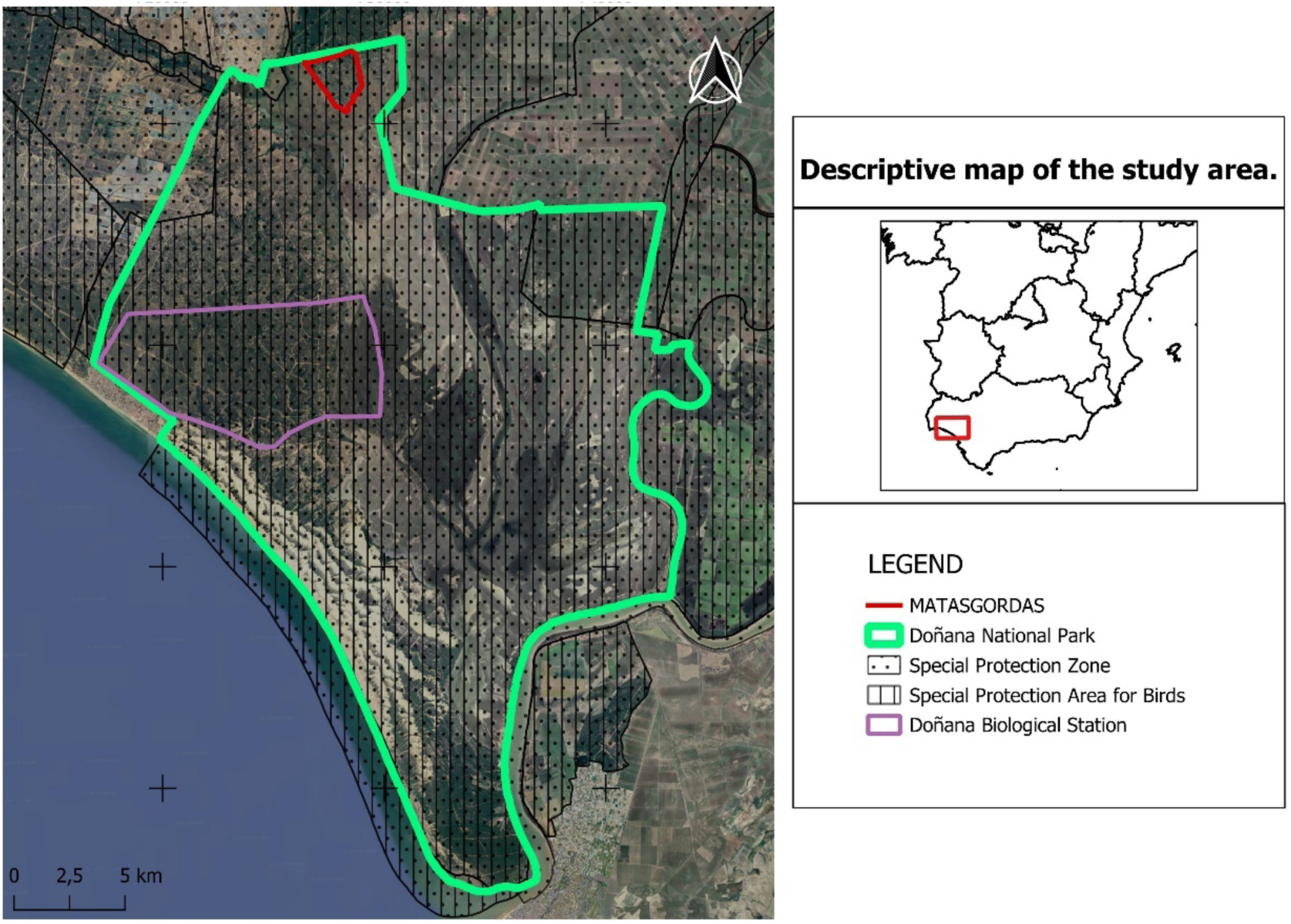
Descriptive map of the study area. The red line indicates the study area, Matasgordas (Google Earth).

The micromammal species present in the study area and respective references are indicated in Table 1 (as for reptiles, see Supporting Information Annex II).

**Table 1.** Micromammal species present in the study area and citations.

| Micromammal species |  | Reference |
| --- | --- | --- |
| Family Muridae |  |  |
| Wood mouse | *Apodemus sylvaticus* | Santoro *et al.*, 2016; Moreno *et al.*, 2009; Palomares & Delibes, 1991; Marfil *et al.*, 2009; Carro *et al.*, 2023; Valverde 1967; Program for Monitoring *Processes and Natural Resources of the Doñana Natural Area*, https://digital.csic.es/handle/10261/287933 |
| Algerian mouse | *Mus spretus* |  |
| Black rat | *Rattus rattus* | Santoro *et al.*, 2016; Carro *et al.*, 2023; Program for Monitoring *Processes and Natural Resources of the Doñana Natural Area*, https://digital.csic.es/handle/10261/287933 |
| Brown rat | *Rattus norvegicus* * | Rojas & Palomo, 2007; Valverde 1967; Program for Monitoring *Processes and Natural Resources of the Doñana Natural Area*, https://digital.csic.es/handle/10261/287933 |
| Family Cricetidae |  |  |
| Southern water vole | *Arvicola sapidus* * | Fedriani *et al.*, 2002; Grilo *et al.*, 2019. Valverde 1967; Program for Monitoring *Processes and Natural Resources of the Doñana Natural Area*, https://digital.csic.es/handle/10261/287933 |
| Mediterranean pine vole | *Microtus duodecimcostatus* * | Grilo *et al.*, 2019; Valverde, 1967; Program for Monitoring *Processes and Natural Resources of the Doñana Natural Area*, , https://digital.csic.es/handle/10261/287933 |
| Family Gliridae |  |  |
| Garden dormouse | *Eliomys quercinus* | Santoro *et al.*, 2016; Moreno *et al.*, 2009; Palomares & Delibes, 1991; Marfil *et al.*, 2009; Carro *et al.*, 2023; Valverde 1967; Program for Monitoring *Processes and Natural Resources of the Doñana Natural Area*, , https://digital.csic.es/handle/10261/287933 |
| Family Soricidae |  |  |
| Greater white-toothed shrew | *Crocidura russula* | Santoro *et al.*, 2016; Moreno *et al.*, 2009; Palomares & Delibes, 1991; Marfil *et al.*, 2009, Carro *et al.*, 2023; Valverde 1967; Program for Monitoring *Processes and Natural Resources of the Doñana Natural Area*, , https://digital.csic.es/handle/10261/287933 |

\*Species that are present in DNP according to the bibliography but are not expected to be found in the study area of Matagordas because it is not their preferred habitat and recent trapping by the Estación Biológica de Doñana indicates their absence.

### 2.2. Camera trap survey

We developed a field protocol which included random systematic sampling in the selected area using CT for the subsequent application of STE model. In parallel, capture-recapture of micromammals was performed using Sherman traps. Both techniques were implemented in late spring and summer 2023.

#### ▪ CTs deployment and settings

Within Matasgordas, we deployed CTs (see Figure 2) in an area overlapping with 4 micromammal monitoring plots (see CR LTERs in Figure 2), where the Biological Station of Doñana (EBD) regularly carries out capture and recapture of micromammals with Sherman traps (see below) for abundance estimation. Thirty CTs were regularly distributed over the study area separated 150 m (Figure 2) from 05/06/2023 to 22/06/2023.

**Figure 2.**
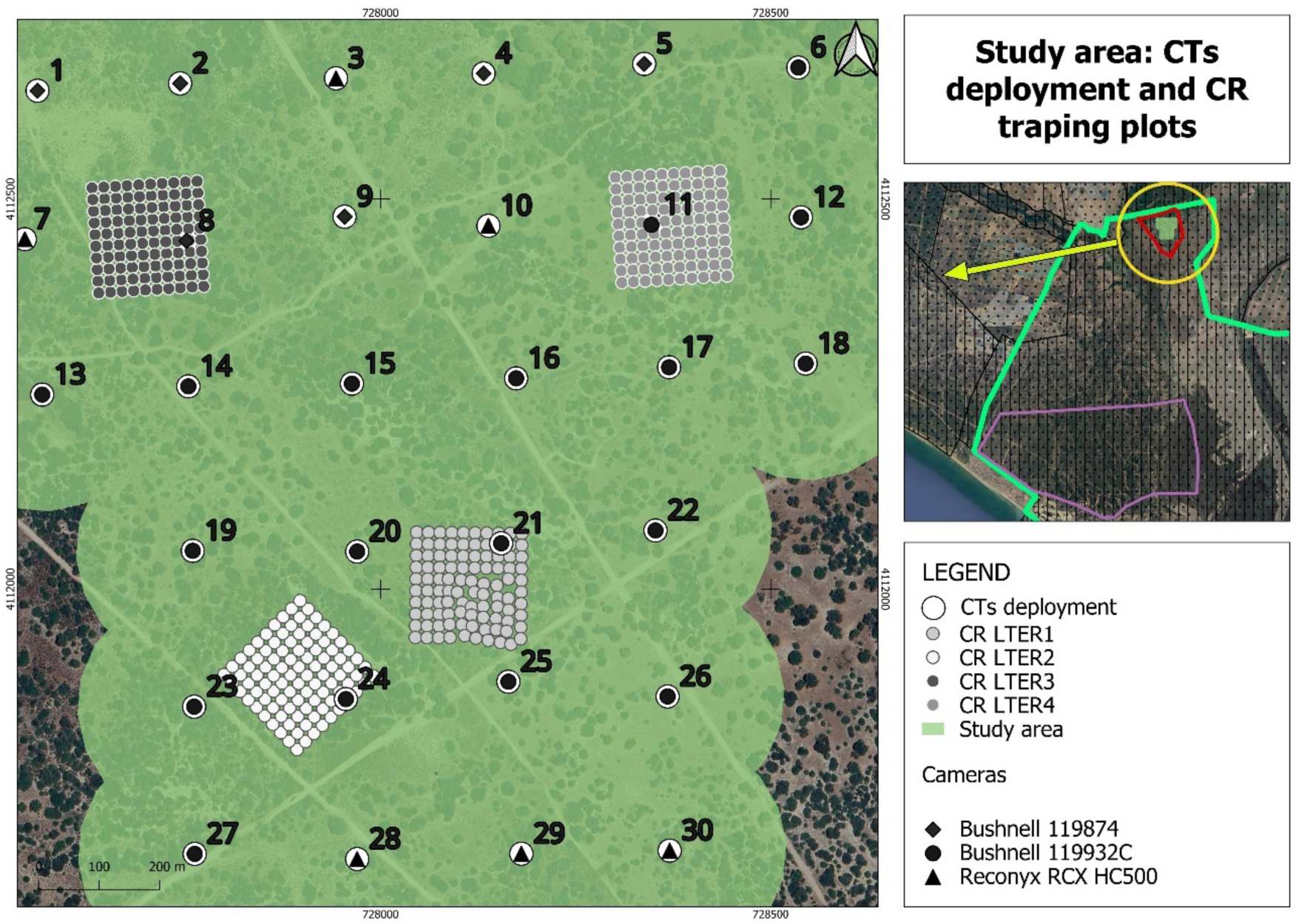
The area where 30 CTs were deployed, indicating the 4 study areas of capture (CR LTER1 to 4) where micromammals were monitored by Sherman traps and Capture-Recapture model. The CT brand is also indicated.

The calculation of micromammal population abundance was achieved through the estimation of the total area sampled by delimiting a 150 m buffer around the coordinates of the CTs, thereby accounting for a total study area of 135.98 ha (see Figure 2). In order to calibrate the area with precision according to the area that is utilised by micromammals (wooded scrubland), a digital photointerpretation was conducted on the orthophoto (National Aerial Orthophoto Plan, PNOA 2022). The normalised difference vegetation index (NDVI) was employed, utilising the near-infrared B8 and red B4 bands of the Sentinel-2 satellite image from 5 February 2023, which was selected due to its cloud-free status. The pixel size employed was 10 metres. Values below 0.2 are predominantly attributed to soil, while values above 0.35 are predominantly attributable to vegetation (Lillesand et al., 2015). By reclassifying the NDVI into two ranges (0.35 value as cut-off), the percentage of shrub and tree vegetation surface was determined to be 36.6% in the study area. Therefore, the calculated study area (woodland) was found to be 49.77 ha once the total area was adjusted by the percentage of shrub and tree vegetation.

The final position of CTs with respect to the planned random systematic sampling coordinates was determined by the availability of microhabitat (covered areas) for micromammals and the expected use by then (given that micromammals typically move under canopy). Therefore, a point within 5-10 metres of the designated location was selected, which was covered by brushes using natural vegetation, or wooden stakes when needed, to attach the CT (Figure 3). CTs were placed at 80 cm (Supporting Information Annex I).

**Figure 3.**
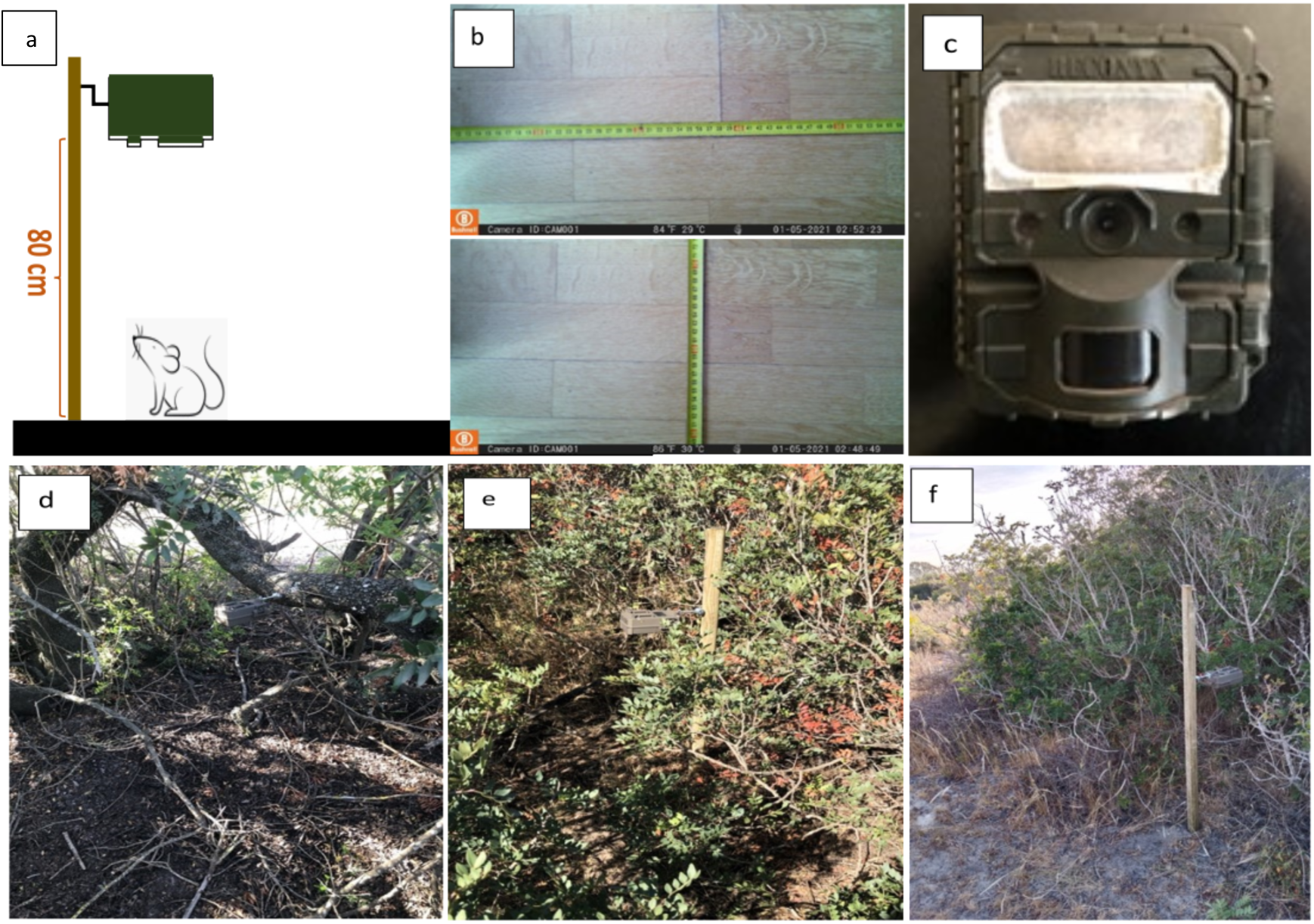
Camera-trap setup. (a) Sketch of deployment, (b) view of the monitoring area for Bushnell model “in the lab”, (c) the infrared led flash brightness is reduced by using semi-transparent white tape; (d,e,f) pictures of real deployments in the field.

Three CT models (Supporting Information Annex I), Reconyx (RCX HC 500) and two Bushnell models (119932C and 119874) were utilised, which allowed for time lapse mode 24 hours, with infrared red light during nocturnal hours. Trigger mode was also operative. The dimensions of the monitored area are 0.13 m^2^ for Bushnell models, and 0.22 m^2^ for Reconyx when placed at 80 cm height. The CTs were programmed to take one picture every minute. The infrared LED flash intensity was reduced with the aid of semi-transparent white tape, as illustrated in Figure 3c. Of the 30 CTs, a total of 25 successfully yielded time-lapse data that was amenable to analysis. Two of the CTs malfunctioned (numbers 29 and 20, Figure 2), and a further three were erroneously set (numbers 1, 2 and 8, Figure 2, which were set only in trigger mode).

##### Capture-Recapture

The density of micromammals was firstly estimated by CR using Sherman traps. The SEMICE protocol (project “Seguimiento de los micromamíferos comunes de España”) was applied with some variations by EBD, as part of the “Program for Monitoring Processes and Natural Resources of the Doñana Natural Area”, with citations available here: https://digital.csic.es/handle/10261/287933. The Sherman traps utilised in this study were equipped with cotton wool inside and food (apples, tenebrionid larvae, and bread soaked in oil). Each of the 4-sampling plots is composed of 100 numbered traps, with an inter-trap distance of 15 metres. Two plots were sampled in May (LTER1 and 2), and two different plots were sampled in July (LTER3 and 4). The traps remained operational for a period of six consecutive nights from 9 May 2023 to 14 May 2023 and were subject to daily inspections in the early morning during May fieldwork. In July, the traps were operational for a period of four consecutive nights from 11 July 2023 to 14 July 2023 and were also subject to daily inspections in the early morning.

The handling was performed in accordance with European legislation (EC Directive 86/609/EEC) and Spanish laws (RD 223/1988; RD 1021/2005), current guidelines for the ethical use of animals in research (ASAB, 2006), the Animal Experiment Committee of CSIC and Government of Andalucía (06/07/59). The handling of captured specimens required the use of latex gloves or gloves made of suitable materials. The rats and dormice were anaesthetised using isoflurane in a custom-built anaesthetic hood of known volume, in which the anaesthetic was administered at a dose of 2-5%. The measurement and weighing of the animals were conducted utilising a 0.5 precision calliper (head-tail, tail, ear and foot length). The animals were marked with an ear-tag (following the disinfection of the material) except for shrews, which were marked with a temporary haircut. After the collection of data, the specimens were conveyed to the original point of capture for immediate release.

### 2.3. Camera trap data analysis

The images were subjected to visualisation on a computer, after which they were annotated with information pertaining to the various species observed (Purroy and Varela, 2016), the date on which the images were captured, and the number of individuals in each species.

#### ▪ Species diversity accumulation curve

The present study evaluated the theoretical effort required to capture all micromammal species present in the area. This evaluation was also conducted for reptiles (Supporting Information Annex II). The function specaccum, which is contained within the vegan package of R (Oksanen et al., 2022), was fitted with the exact method, and the model with the lower AIC value was selected. Subsequently, the curves were plotted using the ggplot package (Oksanen et al., 2024; Wickham, 2016).

#### ▪ Space to event model (STE)

Following Moeller et al. (2021), the application of the model was facilitated by the utilisation of the function ste_built_eh, which is contained within the R package spaceNtime. The total working area was determined by employing the ste_estN_fn function from the specified package.

#### ▪ Capture-recapture (CR)

Due to the limited number of captures and recaptures, we estimated the minimum population density (Castañeda et al., 2018). The calculation of this index considers the number of individuals captured without recaptures within a given area, which in this case is 2.25 ha. Consequently, an estimate of the minimum number of animals that can be found within the specific plots in the study area was obtained.

## 3. Results

### 3.1. Species identification and accumulation curves

The 25 CTs that were ultimately employed for the time-lapse study revealed an average operational lifespan of 12.66 days, with a standard deviation of 6.47 days, ranging from 1 to 17 days. The duration of battery life was a determining factor in the selection of each CT for data collection (see Figure 5). The maximum value of operational days detected was 17 days (CTs 17, 22 and 23), with a median of 16 days. Two CTs malfunctioned, working only 1 day and 7 days, respectively, but the remaining CTs consistently ranged between a narrow window of 15 to 17 days. The time lapse photography was programmed to take one photograph every minute, thereby generating an average of 16,932 images per CT, with the highest value being 22,634 images. A total of 17 different species were detected by CTs (see Table 3), in addition to two taxa/groups of records that remained unidentified at the species level (one out of 478 bird events and one out of 463 reptile events). Of the 17 taxa identified, four were micromammals (*A. sylvaticus, C. russula, E. quercinus* and *R. rattus*), two were lagomorphs, and two ungulates. Three species of birds were detected, with high predominance of *C. ruficollis* (red-necked nightjar). Six reptile species were identified during the study (Supporting Information Annex II). Examples of species detection are illustrated in Figure 4 (for reptiles, see Supporting Information Annex II). In addition to the mentioned species, the capture of *M. monspessulanus* and one unidentified micromammal was also observed in pictures collected by trigger mode, which were not included in Table 2 or the analysis.

**Figure 4.**
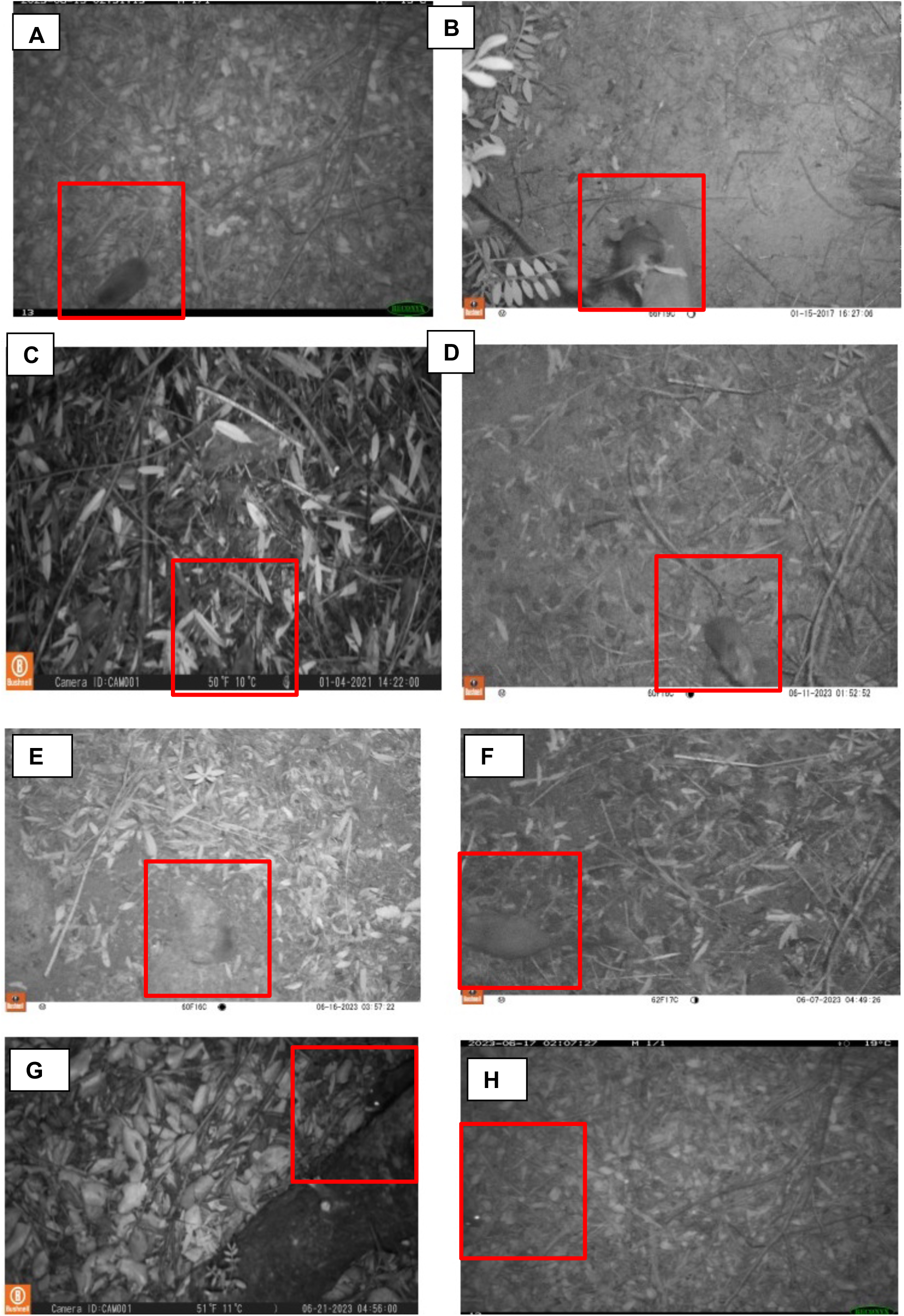
Micromammal captures by camera traps in time-lapse mode. A: *Rattus rattus; B: Elyomys quercinus; C: Crocidura russula; D: Apodemus sylvaticus, E, F, G and H: dubious rodents*.

**Figure 5.**
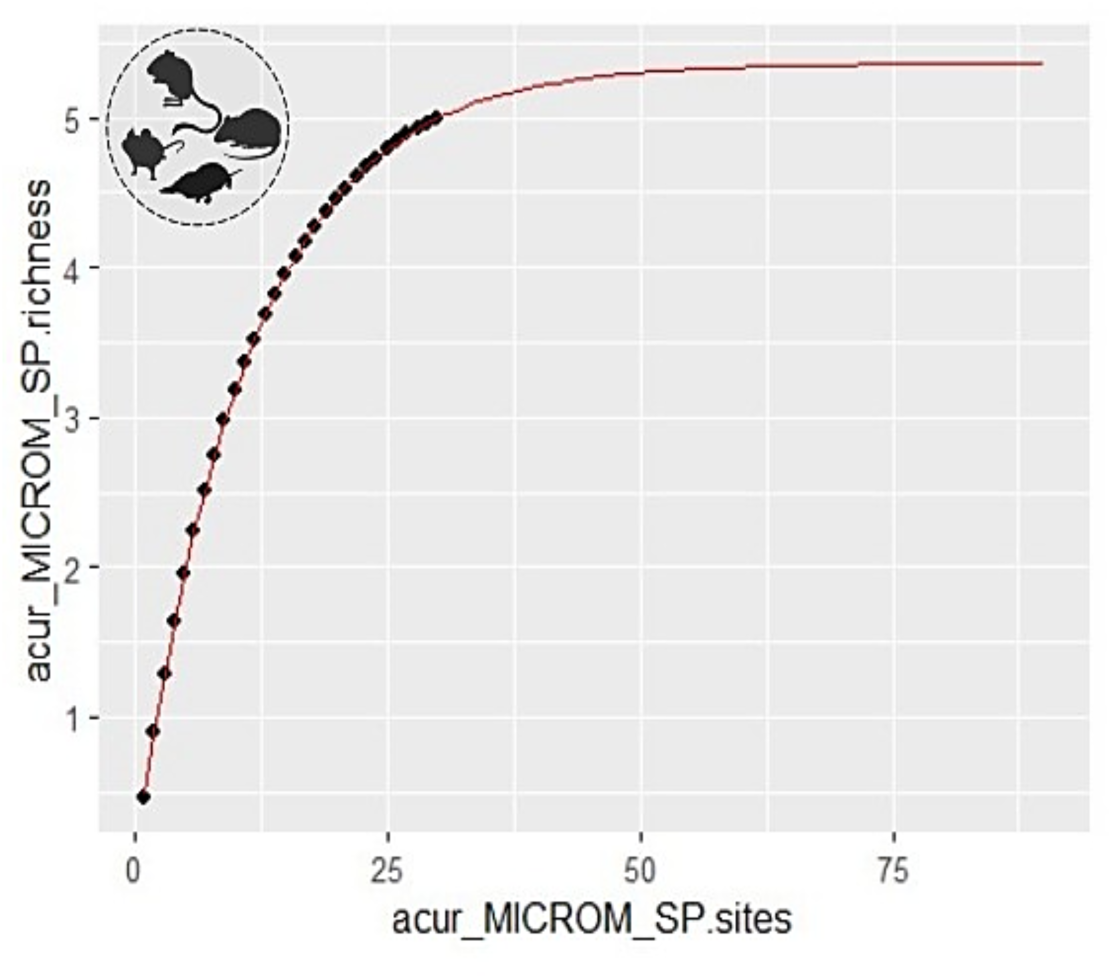
Species accumulation curve for the micromammal species captured in the CTs. Space to event model (STE).

**Table 2.**
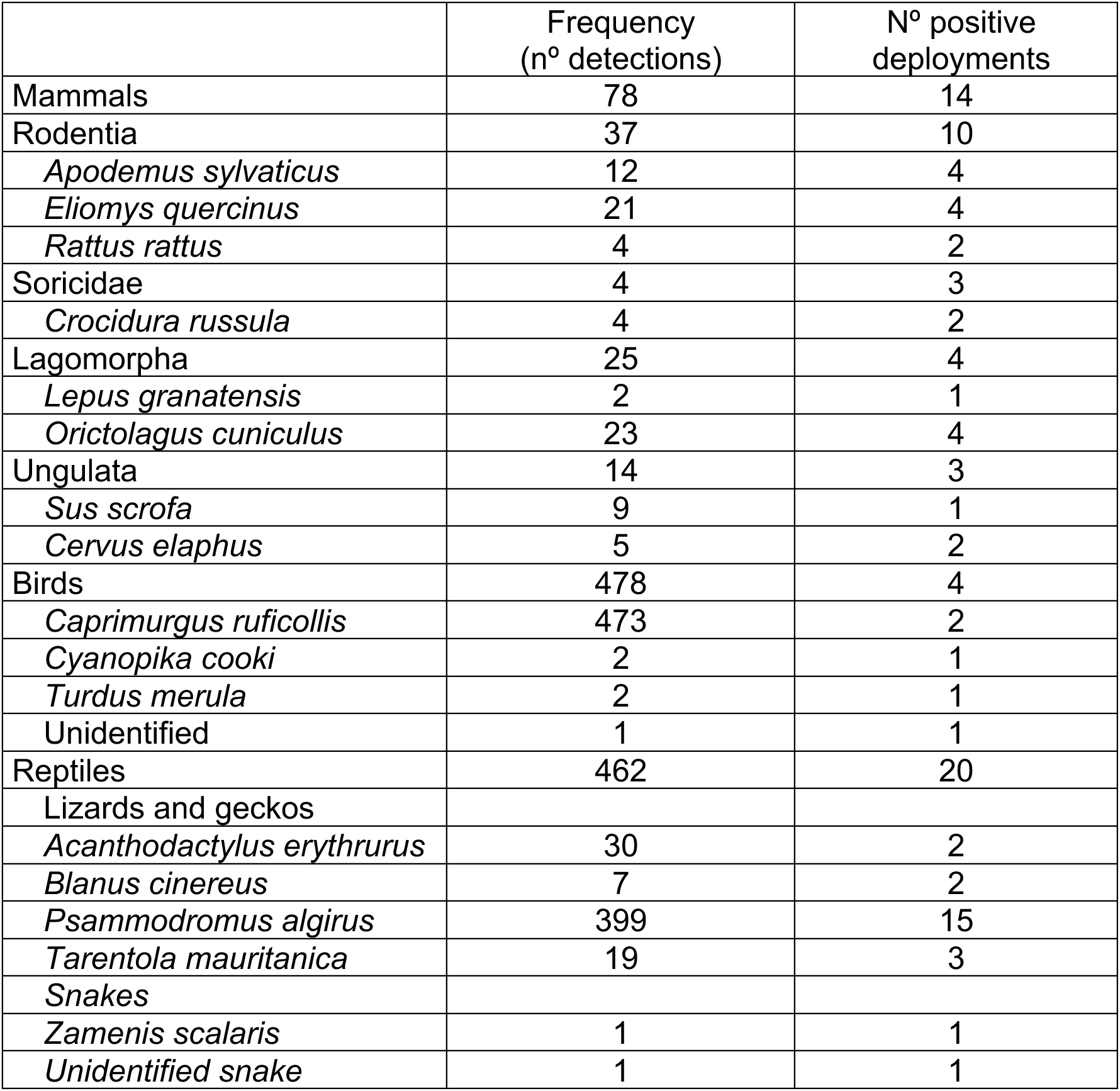
List of identified species and frequency of detection.

**Table 3.** Abundance estimation for micromammal taxa based on STE model. Where N: abundance obtained from the model as number of individuals over 49.77 has study area; SE: standard error obtained from the model; CV: coefficient of variation (expressed as %); Density: density estimated once N is divided by the study area size (ind/ha); LCI-UCI: lower and upper limits for N; CR LTER 1: minimum population (as density, n°/ha) based on captures for plot 1 (LTER 1); CR LTER 2: minimum population based on captures without recaptures for plot 2 (LTER 2); CR LTER 3: minimum population (as density, n°/ha) based on captures for plot 3 (LTER3); CR LTER 4: minimum population based on captures without recaptures for plot 4 (LTER 4).

| SPECIES | N | SE (N) | CV (%) | LCI-UCI (N) | Density CT (N/ha) | Density CR LTER 1 (N/ha) | Density CR LTER 2 (N/ha) | Density CR LTER 3 (N/ha) | Density CR LTER 4 (N/ha) | Average density CR (N/ha) |
| --- | --- | --- | --- | --- | --- | --- | --- | --- | --- | --- |
| *Apodemus sylvaticus* | **26.64** | 13.28 | 49.83 | 10.58-67.067 | **0.54** | **1.88** | **2.33** | **0** | **0** | 1.05 |
| *Crocidura russula* | **13.53** | 9.46 | 69.90 | 3.92-46.56 | **0.27** | 0 | 0 | **0.78** | **0** | 0.20 |
| *Elyomys quercinus* | **39.49** | 16.16 | 40.91 | 18.26-85.38 | **0.79** | **1.88** | **0.39** | **0.39** | **0.78** | 0.86 |
| *Rattus rattus* | **13.55** | 9.46 | 69.84 | 3.93-46.60 | **0.27** | 0 | 0 | 0 | **0.39** | 0.10 |
| **Micromammals (Total)** | **66.34** | 20.94 | 31.57 | 36.26-121.37 | **1.33** | 3.96 | 2.72 | 1.17 | 1.17 | 2.26 |

The groups with the highest detection frequencies were birds (46.95%) and reptiles (45.38%), followed by mammals (7.66%), of which 4.03% were micromammals and 3.63% were medium- and large-sized mammals).

The accumulation curve of micromammal species (Figure 5) indicated a theoretical maximum expected species number of 5.36 (which coincides with the five most frequently caught species in the area according to monitoring data and literature), compared with the four detected by CTs, and the eight that are expected to be present in DNP beyond Matasgordas. *M. Spretus* and *M. duodecimcostatus* were not detected in the study area. Similarly, *A. sapidus* and *R. norvergicus* were not detected. This would represent 74.58% of the species present in Matasgordas, with an effort of 28 CTs (accounting for an average of 12.7 CT*days operational time). This elementary analysis demonstrated that a sample size of fewer than 40 CTs (given the duration of this study) for the applied period would be adequate to capture the five predicted micromammal species. The species accumulation curve for the reptilian species captured in the CTs is presented in Supporting Information Annex II.

The micromammal species densities ranged from 0.27 to 0.79 individuals per hectare (Table 3). The lowest values were recorded for *C. russula* and *R. rattus* (0.27 ind/ha each). The group of micromammals did not reach even half of the values for reptile species (Table 3, Figure 6 and Supporting Information Annex II). The reptile group has higher abundance than the micromammal group, with values of 38.93 individuals/hectare compared to 1.33 individuals/hectare.

**Figure 6.**
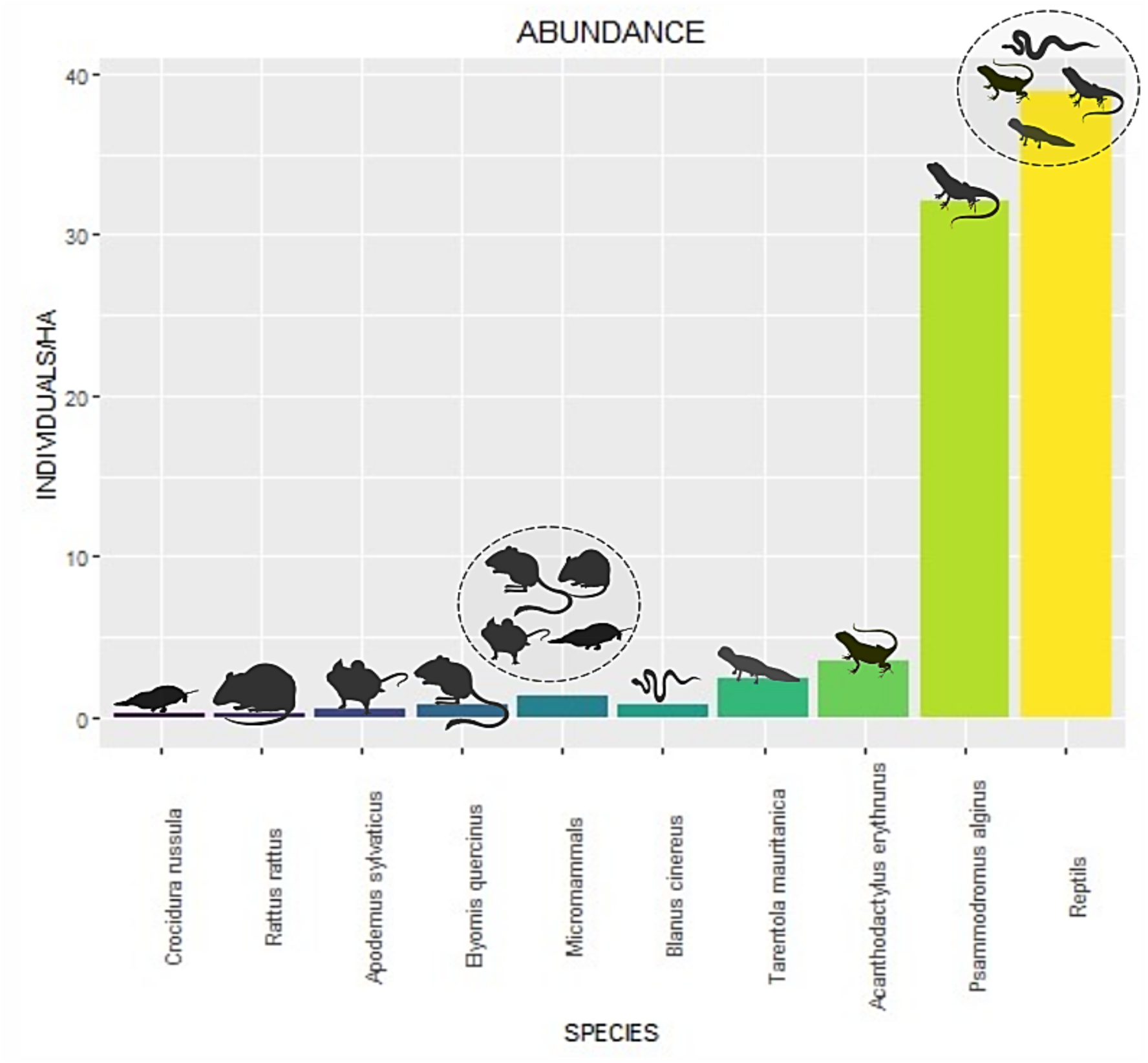
Abundance of different species and groups obtained by the STE model.

#### Capture-recapture (CR)

Given the low number of micromammal recaptures (Table 4), the capture-recapture model was not applied, and a minimum population abundance was instead estimated. A total of two species were captured during the May trapping: *E. quercinus* and *A. sylvaticus*. Furthermore, during the July trapping, in addition, *R. rattus* and *C. russula* were also captured. The values obtained from each plot are presented in the following section, alongside the estimated abundances for each plot (Tables 3 and 4).

**Table 4.** Results of capture-recapture of micromammals.

| Plots | Nº captures of different individuals | Minimum density | Capture rate | Recaptures |
| --- | --- | --- | --- | --- |
| LTER1 | 6 *E. quercinus* | 2.66 ind/ha | 1% | 1 |
| LTER1 | 6 *A. sylvaticus* | 2.66 ind/ha | 1% | 5 |
| LTER2 | 2 *E. quercinus* | 0.88 ind/ha | 0.33% | 2 |
| LTER2 | 13 *A. sylvaticus* | 5.77 ind/ha | 2.16% | 4 |
| LTER3 | 2 *E. quercinus* | 0.88 ind/ha | 0.5% | 2 |
| LTER3 | 4 *C. russula* | 1.77 ind/ha | 1% |  |
| LTER4 | 2 *R. rattus* | 0.88 ind/ha | 0.5% |  |
| LTER4 | 4 *E. quercinus* | 1.77 ind/ha | 1% | 1 |

The species with the highest abundance was *A. sylvaticus*, reaching minimum population densities of 5.77 individuals/ha in plot 2. For *E. quercinus*, the values were lower, except for plot 1, which presented similar values. *C. russula* was present only in LTER 3, with 1.77 ind/ha, which is higher than the 0.88 ind/ha recorded for *E. quercinus*. Finally, LTER 4 was the only plot where *R. rattus* was present, with an average of 0.88 ind/ha.

#### Comparison between time lapse camera-trapping and capture

The application of time lapse and capture techniques yielded analogous results with regard to the diversity of micromammal species (Figure 7) and there was a high degree of concordance.

**Figure 7.**
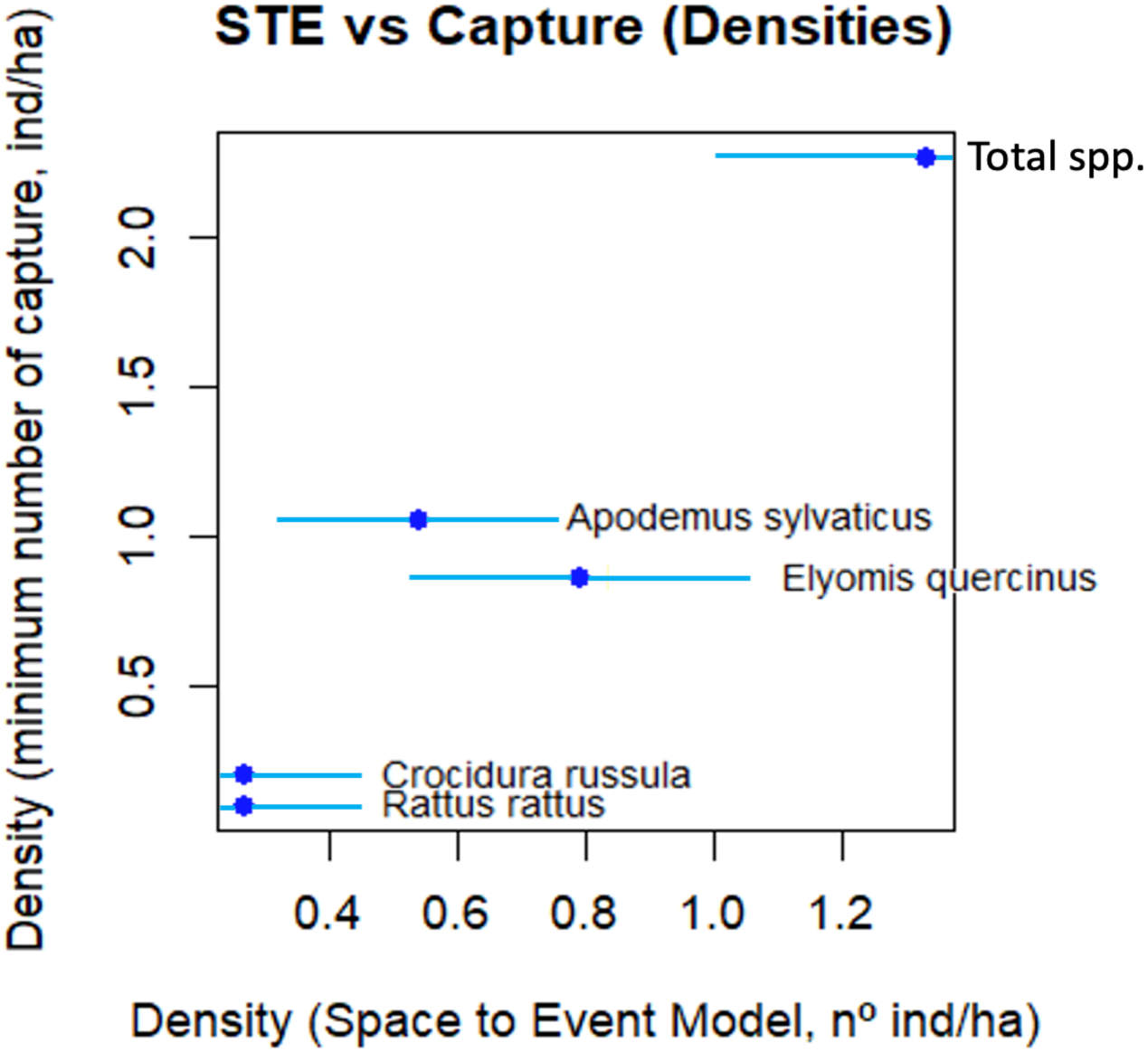
Agreement between the abundances obtained with Space to Event Model (X-axis) *versus* capture (Y-axis, average value for the 4 plots) for different species of micromammals. The horizontal bars indicate SE.

### 3.2. Effort and practical issues

A total of 492,691 time lapse images (from 25 CTs in time lapse mode) were utilised for density analysis, of which 1,018 images corresponded to detections. Following a thorough evaluation of the efforts applied, it was determined that both CT and CR required similar levels of involvement. CT generated an effort of 309.6 hours per person, while CR required a marginally higher effort, totalling 360 hours per person (Table 5).

**Table 5.**
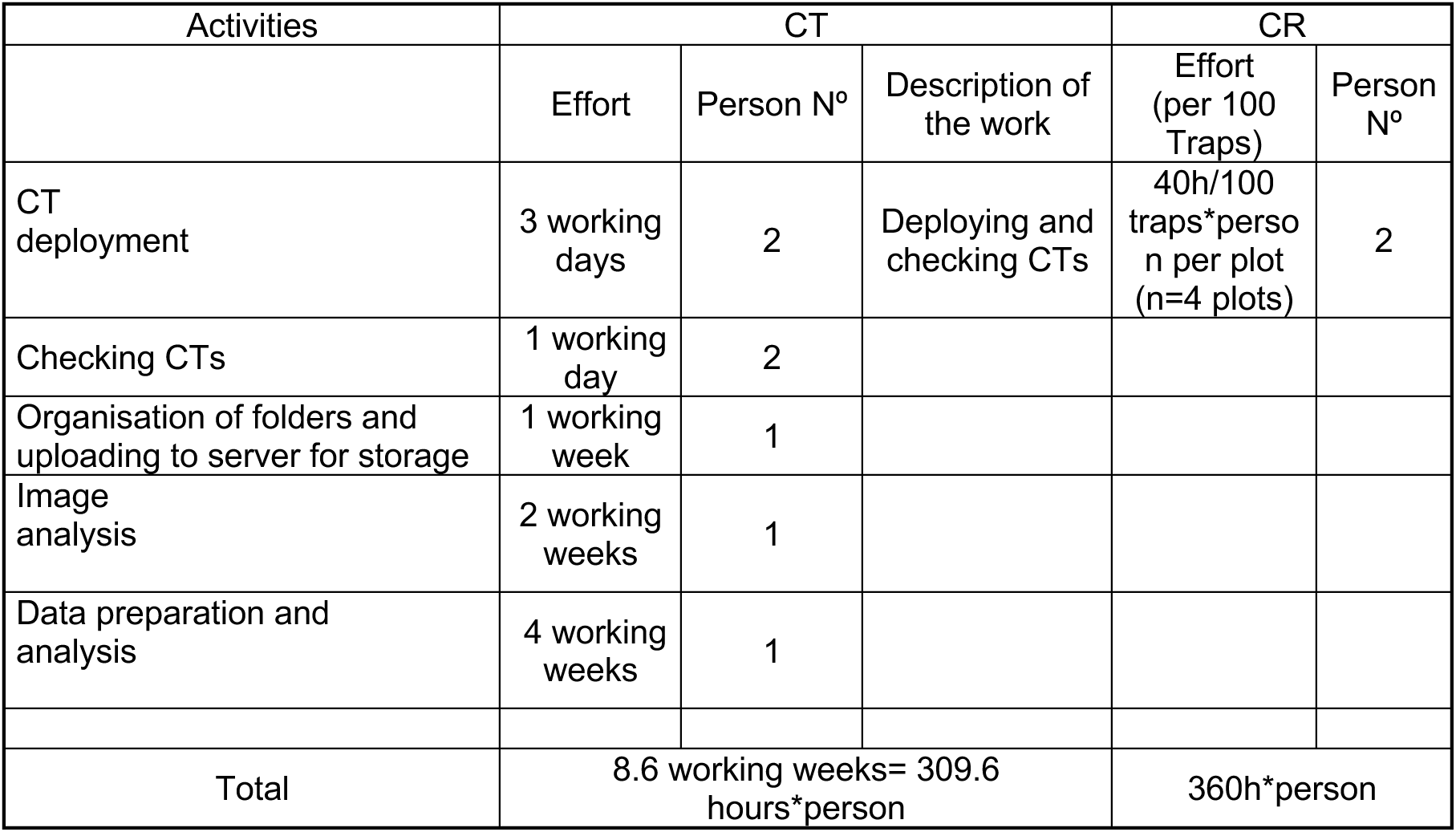
Breakdown of efforts spent on CT and CR methodologies.

| Activities | CT |  |  | CR |  |
| --- | --- | --- | --- | --- | --- |
|  | Effort | Person N° | Description of the work | Effort (per 100 Traps) | Person N° |
| CT deployment | 3 working days | 2 | Deploying and checking CTs | 40h/100 traps*person per plot (n=4 plots) | 2 |
| Checking CTs | 1 working day | 2 |  |  |  |
| Organisation of folders and uploading to server for storage | 1 working week | 1 |  |  |  |
| Image analysis | 2 working weeks | 1 |  |  |  |
| Data preparation and analysis | 4 working weeks | 1 |  |  |  |
| Total | 8.6 working weeks= 309.6 hours*person |  |  | 360h*person |  |

## 4. Discussion

This study was the first, to the best of our knowledge, to test a time-lapse based camera-trapping protocol and the application of STE to determine the abundance of micromammals without the need to capture or handle them. The species richness and densities of micromammals present in the area were found to be well-correlated with the methodology of capture. We evidenced the utility of this approach for the monitoring of other small-sized terrestrial wildlife (incl. reptiles) and its potential for CT multi-species monitoring. In terms of practicability, field and data processing efforts are both deemed feasible and will continue to improve as new automatic identification tools are incorporated.

### 4.1. Practical issues of camera trap deployment

The mean value of CT operational days averaged almost 13 days, which was consistent over most CTs (between 15-17 days). This can be attributed to the battery life of the Alkaline AA batteries, which were depleted before the CT memory reached its maximum capacity. In order to increase the operational life without the necessity of changing batteries and maintaining the same settings, the following measures can be taken: i) higher power batteries can be used, (e. g., lithium), ii) the trigger mode can be deactivated, ¡ iii) external batteries and/or portable solar panels can be used, and the intensity of the infrared flash can be reduced (Iv). In case of a reptile-focused approach, the operational period of CTs can be set only for daytime. Following the favourable outcomes of the initial approach, we believe that CT models can be developed specifically for time-lapse applications, thereby eliminating the necessity for PIR sensors.

Our results suggest that augmenting the sampling effort, for example by extending the operational time of CTs for our given number of CTs, would likely enhance the precision of estimations and species richness determination. Further research is required to evaluate effort in terms of the number of CTs and the duration of monitoring. Additionally, the higher frequency of time-lapse would likely contribute to a reduction in the duration of field sampling (Kays *et al.,* 2020). This is a particularly pertinent consideration when the necessity arises to undertake a rapid assessment of wildlife populations.

The focal distance to the ground was adapted to the characteristics of the CT models employed in this study (at 80 cm). However, it is possible to improve the quality of the image focus by using lenses to focus on small objects, facilitating the acquisition of higher-quality images and more detailed information for species identification (Nardotto and Bertolin, 2024).

### 4.2. Species identification and richness

The species of micromammals that appear most frequently in the study area according to recent literature are *A. sylvaticus*, *C. russula*, *E. quercinus*, *M. spretus* and *R. rattus* (Carro et al., 2021; Santoro et al., 2016). Of these five species, *M. spretus* is the only one that has not been captured either by Sherman traps or by CT. Torre et al. (2023) posit that the presence of *M. spretus* is negatively correlated with vegetation complexity and water deficit. Furthermore, the recruitment of juveniles is reduced in forested landscapes. The critical water situation in DNP (Green et al., 2024) may be exerting a negative influence on the population of the species, and consequently on its detectability, specifically in Matasgordas (Smith et al., 2023).

As indicated in the DNP, there may be some species present in the area which were not captured by either method (*A. sapidus*, *M. duodecimcostatus* and *R. norvegicus*), which was expected since the study area does not coincide with their preferred habitats. The first species has been observed to demonstrate a preference for riverside habitats and areas near seasonal ponds within the DNP (Mate et al., 2013; Román, 2003). Conversely, *R. novergicus* has been found to select rural and urban environments, sewers, and rubbish dumps (Rojas and Palomo, 2007). In the specific case of *M. duodecimcostatus*, its subterranean habits may be related to its absence in camera trapping and captures (Cotilla and Palomo, 2007). This highlights the necessity for the implementation of specific protocols for the determination of density, particularly in the context of subterranean species.

One of the most challenging aspects of image analysis pertains to the identification of small wildlife, such as micromammals, which are also hardly ever observed in daylight hours. The employment of specific distinguishing features in black-and-white, such as the disproportionately large size of the eyes and ears relative to the head and the significantly elongated tail relative to the head and body were hallmarks of the species under study (Purroy and Varela, 2016; Turón Artigas, 2021). The implementation of this approach demanded the expertise of the research team.

The performance of our CT protocol was satisfactory, as indicated by the species accumulation curve. This curve is a relevant tool for the comparison of small terrestrial species diversity within specified communities and for the determination of the necessary effort required to describe the diversity of such communities (Gotelli and Colwell, 2001).

### 4.3. Abundance

#### ▪ Space to Event Model

The estimated densities of micromammals obtained have been low, as expected. The most abundant species was *E. quercinus* (0.79 ind/ha), followed by *A. sylvaticus*, *R. rattus* and *C. russula*, giving a total value of 1.33 ind/ha for all micromammal species. Moreno et al. (2016) reported the virtual disappearance of *E. quercinus* and *R. rattus*, a marked decrease in *A. sylvaticus*, and the dominance of *M. spretus*. However, the latter species was not captured throughout the study. Moreno et al. (2009) cited *M. spretus* as the dominant species, while *E. quercinus* showed a very low abundance. *A. sylvaticus* exhibited spatial avoidance mechanisms in the presence of *M. spretus*. The absence of *M. spretus* may be attributable to its predilection for open environments, while *A. sylvaticus* demonstrates a preference for areas covered by vegetation (Santoro et al., 2016). Specifically, in the case of *A. sylvaticus*, the 0.54 ind/ha recorded in summer 2024 are lower than the 2.5 ind/h determined by Jubete (2007) from May to July, indicating an abundance 5 times lower. *A. sylvaticus* demonstrates a marked preference for wetter environments in summer, such as heathlands (Torre et al., 2002). Acorns from cork oak trees represent a vital food source for *Apodemus*, and the decline of cork oak trees and the severe drought may also be a determining factor in the population dynamics of the species (Docampo et al., 2019). *E. quercinus* is the species with the highest abundance; however, it is still far from those cited by Moreno (2007) in 1980-1984, where up to 30 ind/ha were counted. The abundance of micromammals in DNP was once one of the highest in the Mediterranean region (Moreno et al., 2009). Docampo et al. (2019) expressed concerns regarding the conservation of DNP ecosystems, citing the rapid and extensive morphological changes observed in *A. sylvaticus*.

The present study demonstrated the feasibility of conducting multispecies monitoring for small wildlife through the utilisation of CTs in time-lapse mode, followed by the implementation of STE. This phenomenon is of particular concern in the Mediterranean region, where the decline of keystone species, such as rabbits, and the concomitant increase in wild ungulate populations may have had direct consequences for Mediterranean communities and associated ecosystems (see Moreno et al., 2007; Muñoz et al., 2009; Carpio et al., 2014; Delibes-Mateos et al., 2009). Whilst there is evidence to suggest that the community of micromammals in DNP is undergoing change, further research is required to improve our protocol. This should include the deployment of CT in open sites, in addition to the current CT disposal in covered areas. Furthermore, it is necessary to increase the level of effort expended.

#### ▪ Comparison between Time to Event Model and direct capture for micromammals

It has been determined by several studies that CT surveys of small terrestrial mammals may provide a cost-effective technique for surveying small terrestrial mammals, with results that are comparable to those of live trapping (e.g., De Bondi et al., 2010). However, no time-lapse approach has been tested, and thus the added value of providing density and precise quantitative configuration of the community remains to be ascertained. The employment of time lapse, in conjunction with the subsequent implementation of the STE, and the utilisation of direct capture, as evidenced in this particular instance, has yielded analogous outcomes with respect to micromammal species richness. The species accumulation curves of CT, as previously discussed, have proven to be satisfactory; nevertheless, they are amenable to enhancement through the augmentation of the sampling effort.

### 4.4. Effort and practical issues

CTs and STE did not result in a reduction in efforts when compared to CR. However, field and data processing efforts are both feasible and will continue to improve as new automatic identification tools are incorporated into the protocol (the images were analysed manually) and several aspects of the protocol are improved. Furthermore, CTs are not invasive, and did not give rise to ethical issues or staff safety concerns, while concomitantly expanding the range of monitored taxa. The utilisation of artificial intelligence tools, which employ machine learning and computer analysis to expedite the automation of image labelling, has the potential to diminish analysis effort and is becoming increasingly prevalent (Leorna and Brinkman, 2024). Firstly, to automatically filter blank images (which constitute the vast majority in time lapse mode) from images featuring wildlife, and, secondly, to classify of species with increasing sensitivity (Veléz et al., 2022). Whilst these methodologies have the potential to engender time and effort savings, powerful computational systems may be required when dealing with large quantities of photographs.

The following conclusions were drawn: firstly, the utility of the approach for monitoring other small terrestrial wildlife, such as reptiles, was evidenced; secondly, the potential for the approach to contribute to CT multi-species monitoring was demonstrated; thirdly, the practicability of the approach was evidenced; and finally, it was demonstrated that field and data processing efforts were feasible and will only improve as new tools are used.

## Supporting information

Supporting information on reptiles

## Acknowledgements

We would like to thank technicians at the Doñana Biological Station, all the trainees who have worked with us, and the Doñana National Park Staff. This is a contribution of Biodiversa+ project “Big_Picture” (ref. Proyecto PCI2024-153504 convocatoria europea Biodiversa+, funded by MICIU/AEI) and Plan Nacional ref. PID2022-142919OB-100.

## Authors’ Contributions Statement

AE, JV, FC and PA conceived the ideas and designed methodology; AE, FC and VGL collected the data; AE, JV and VGL analysed the data; AE, JV and VGL led the writing of the manuscript. All authors contributed critically to the drafts and gave final approval for publication

