## Supporting information on reptiles for "A look at the ground: time lapse camera trapping to determine the composition and abundance of small sized wildlife communities"

**Annex I.** Table 1A. Camera trap settings for different models

| Set mode | Camera-trap Model and Setting (Bushnell Trail Cameras) | | |
|---|---|---|---|
| | 119922C | 119874 | 119874 |
| Mode | Camera | Camera | Camera |
| Interval | 1 s | 0.6 s | 1 s |
| Image Size | 3 MP | HD | 8 MP |
| Image Format | NA | Full Screen | Full Screen |
| Capture Number | 1 Photo | 3 Photos | 3 Photos |
| Led Control | NA | Low | Low |
| Additional Settings | On | NA | NA |
| Sensor Level | Normal | Normal | Normal |
| NV Shutter | NA | High | High |
| Field Scan | On | On | On |
| FS Interval | | | |
| FS Periods (A) | Start: 12:00 Stop: 23:59 | Start: 12:00 Stop: 23:59 | Start: 12:00 Stop: 23:59 |
| FS Periods (B) | Start: 00:00 Stop: 11:59 | Start: 00:00 Stop: 11:59 | Start: 00:00 Stop: 11:59 |
| Flash Mode | Fast Motion | NA | NA |
| Time Stamp | On | On | On |
| Camera Mode | 24 Hours | 24 Hours | 24 Hours |
| Set Clock | Local time | Local time | Local time |
| **Set Mode** | **Camera-trap Model and Setting (Bushnell Trail Cameras)** | | |
| | RCX HP2 | RCX HC500 | |
| Motion Pictures | On | On | |
| Number of Pics | 3 | 3 | |
| Time Btw Pic | Rapidfire | Rapidfire | |
| Motion Video | Off | Off | |
| Quiet Period | No Delay | No Delay | |
| Sensitivity | Medium | Medium | |
| Time Lapse | On | On | |
| Lapse Video | Off | Off | |
| Interval | 1 min | 1 min | |
| Lapse Schedules | 24 Hour | 24 Hour | |
| Take Pictures | Day/Night | Day/Night | |
| Flash Output | Low | Low | |
| Night Mode | Fast Shutter | Fast Shutter | |

**Annex II**. This Annex presents the reptile species that are present in DNP (Table 1b), results obtained on these taxa in the present paper, and the main considerations regarding the use of STE to determine reptile abundance.

Regarding the reptile group, the species that are present in DNP are presented in Table 1b.

Table 1b. Reptile species present in the study area (source: *Program for Monitoring Processes and Natural Resources of the Doñana Natural Area*, and citations there in, https://digital.csic.es/handle/10261/287933 and citations in table). We have been included the species associated with Mediterranean scrub and pine forest areas in DNP.

| Reptile species | | Reference |
|---|---|---|
| Family Blanidae | | |
| *Blanus cinereus* | Blind shingles | Valverde, 1967; Paniagua & Rivas, 1987 |
| Family Chamaleonidae | | |
| *Chamaleo chamaleon* | Common chameleon | Andreu, 2014; Rodríguez, 2002 |
| Family Colubridae | | |
| *Coronella girondica* | Southern smooth snake | Andreu, 2014; Rodríguez, 2002 |
| *Hemorrhois hippocrepis* | Horseshoe snake | Andreu, 2014, Rodríguez, 2002 |
| *Macroprotodon brevis* | Western Garter Snake | Andreu, 2014; Rodríguez, 2002 |
| *Natrix maura* | Water snake | Valverde, 1967; Paniagua & Rivas, 1987 |
| *Natrix natrix* | Collared snake | Andreu, 2014; Rodríguez, 2002 |
| *Rhinechis scalaris* | Ladder snake | Andreu, 2014; Rodríguez, 2002 |
| Family Lacertidae | | |
| *Acanthodactylus erythrurus* | Red-tailed lizard | Valverde, 1967; Arribas, 2023 |
| *Timon lepidus* | Ocellated lizard | Valverde, 1967; Arribas, 2023 |
| *Psammodromus algirus* | Long-tailed lizard | Valverde, 1967; Arribas, 2023 |
| *Psammodromus occidentalis* | Iberian western nettle | Valverde, 1967; Arribas, 2023 |

| | | |
|---|---|---|
| *Podarcis carbonelli* | Carbonell´s lizard | Andreu, 2014; Rodríguez, 2002 |
| Family Lamprophophiidae | | |
| *Malpolon monpessulanus* | Western bastard snake | Andreu, 2014; Rodríguez, 2002 |
| Family Phyllodactylidae | | |
| *Tarentola mauritanica* | Gecko | Valverde, 1967; Arribas, 2023 |
| Family Scincidae | | |
| *Chalcides bedriagai* | Iberian sling | Rodríguez, 2002 |
| *Chalcides striatus* | Tridactyl sling | Rodríguez, 2002 |
| Family viperidae | | |
| *Vipera latastei* | Snout-nosed viper | Andreu, 2014; Rodríguez, 2002 |

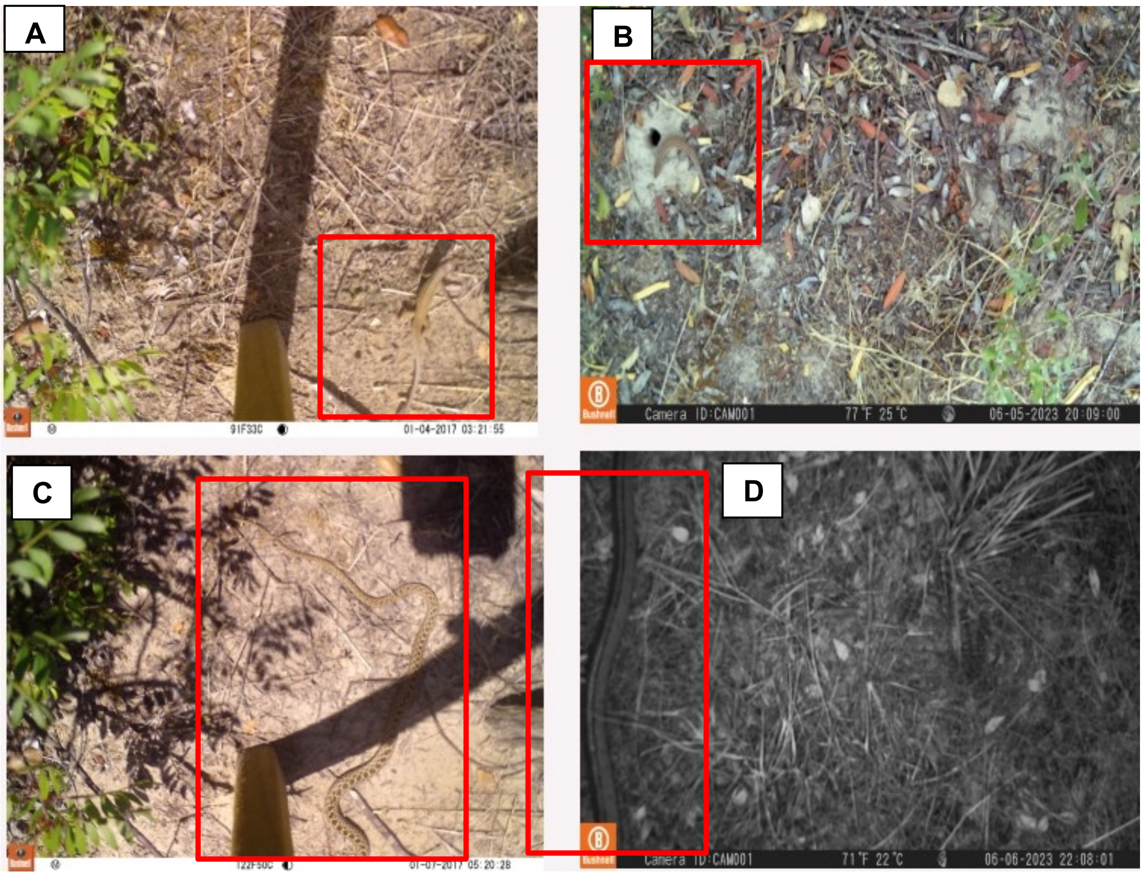

Figure 1b. Examples of reptiles captured by the CTs. A: *Psammodromus algirus; B Acanthodactylus erythrurus; C: Malpolon monspessulanus; D: Zamenis scalaris.*

With regard to the species accumulation curve of reptiles, a predicted value of 8 species in the study area was obtained. The six captured species would theoretically contribute to a total of 71.89% of the reptile species richness in the study area of Matasgordas. Of the species expected to be found in the area (see Table 1b), the ocellated lizard (*T. lepidus*), the Bedriaga`s skink (*C. bedriagai*), and the water snake (*N. maura*) were not captured by time lapse. With regard to the sampling effort for reptiles, it is imperative to exceed a value of 50 trapping CTs (given the duration of this study) to reach the asymptote of species diversity detection, in comparison to the less than 40 that would be necessary for micromammals (see Figure 2b, left).

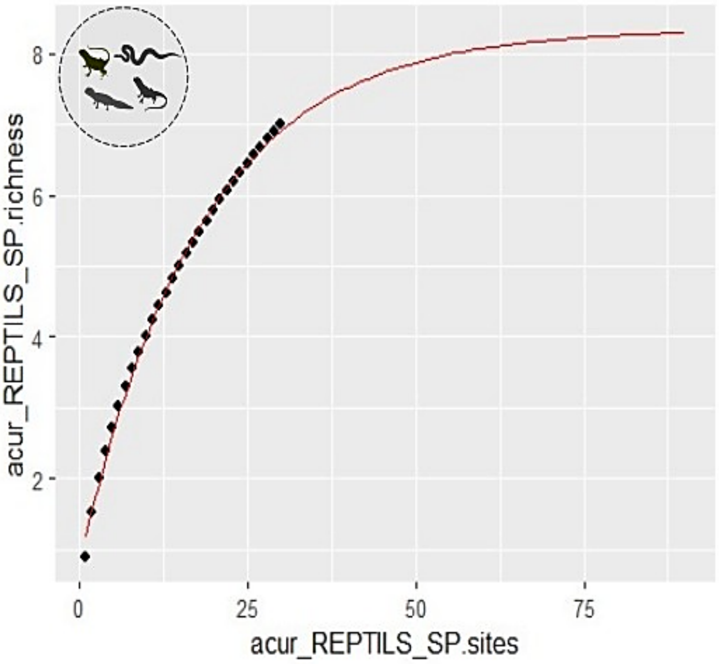

Figure 2b. Species accumulation curve with the reptile species captured in the CTs.

The species with the highest abundance was, by far, *P. algirus*, with an average of 32.08 individuals per hectare (ind/ha). The lowest values were recorded for *C. russula* and *R. rattus* (0.27 ind/ha each). The reptile group has higher abundance than the micromammal group, with values of 38.93 individuals/hectare compared to 1.33 individuals/hectare (see Table 2b and Figure 2b). The species with the highest abundances after *P. algirus* were also reptiles: *A. erthrunus* and *T. mauritanica*. Nevertheless, the group of micromammals did not attain even half of the values recorded for the specified reptile species (see Figure 2b).

Table 2b. Abundance estimation for reptile taxa based on STE model. Where N: abundance obtained from the model as number of individuals over 49.77 has study area; SE: standard error obtained from the model; CV: coefficient of variation (expressed as %); Density: density estimated once N is divided by the study area size (ind/ha); LCI-UCI: lower and upper limits for N.

| SPECIES | N | SE (N) | CV (%) | LCI-UCI (N) | Density CT (N/ha) |
|---|---|---|---|---|---|
| *Acantodactylus erythrurus* | **179.57** | 34.46 | 19.90 | 123.7-260.67 | **3.61** |
| *Blanus cinereus* | **39.82** | 16.22 | 40.74 | 18.47-85.84 | **0.80** |
| *Psammodromus algirus* | **1596.58** | 102.94 | 6.45 | 1407.22-1811.41 | **32.08** |
| *Tarentola mauritanica* | **119.66** | 28.13 | 23.51 | 75.95-188.51 | **2.40** |
| ***Reptiles (Total)*** | **1937.33** | 113.45 | 5.86 | 1727.46-2172.68 | **38.93** |

*Main considerations regarding the use of STE to determine reptile richness and abundance*

A wide range of terrestrial reptiles have been reported in DNP, with some species exhibiting a marked preference for Mediterranean scrub. The species accumulation curve predicts a value of 8 species for an effort of 90 CTs (30TCs were deployed), while 6 captured species were detected in the Matasgordas study area. With regard to the issue of absences within the context of our sampling, it is important to note that the study area may not coincide with the preferred habitat of *P. occidentalis* (although this species was detected once in trigger mode), as it has a documented preference in Huelva for areas characterised by minimal plant cover (Fietze, 2012). *P. carbonell* thrives in areas with high densities of scrub, being more prevalent in areas that have undergone thinning (Roman *et al.,* 2006). Ocellated lizards (*T. lepidus*) have experienced a significant decline in population numbers in DNP (Andreu, 2014), likely attributable to their role as alternative prey to declining rabbit populations (Salgado & Hernández, 2013). With regard to the identification of species, the same difficulty arises as in the case of micromammals, since occasionally, the images obtained were not clear.

As stated in the 2023 Report on the State of Biodiversity in DNP (Arribas, 2023), the Mediterranean scrub has been found to contain a greater number of reptile species than the dunes. This report indicates that the most prevalent species was the long-tailed lizard (*P. algirus*), followed by the common gecko (*T. mauritanica*) and the red-tailed lizard (*A. erythrurus*), among other species (Arribas, 2023). Despite the utilisation of a divergent methodology in comparison with the present study, and the consequent unfeasibility of a comprehensive comparison, it is noteworthy that *P. algirus* exerts a dominant influence over the other two species, as evidenced in the present study. Mellado (1980) posited that adults are primarily associated with dense scrubland formations, while juveniles demonstrate a greater propensity for utilising a more diverse array of resources. This could imply a greater probability of capture. The Atlas and Red Book of Amphibians and Reptiles of Spain (Carretero *et al*., 2002) indicates that this species reaches high densities when shrub cover is significant, which coincides with the geographical location of our study area. In the case of *A. erythurus*, it has been recorded that in DNP it makes a more restricted use of space in open spaces, where the substrate present is loose sand, given the morphology of its toes (Mellado, 1980). In order to ascertain the reliability of the range of estimated abundances and thereby determine their usefulness for monitoring purposes, a comparison with other sampling techniques, such as distance sampling transects, is desirable in the future. Isolated populations of lizards have been identified as being particularly vulnerable to extinction, which implies their importance from a conservation point of view (Carretero et al., 2008). It is imperative to devise pragmatic monitoring programmes that can be implemented within the framework of multi-taxa monitoring. Evidence has demonstrated that CTs are a promising, albeit untapped, tool that requires further refinement.
